# Optical properties of living corals determined with diffuse reflectance spectroscopy

**DOI:** 10.1101/563106

**Authors:** Steven L. Jacques, Daniel Wangpraseurt, Michael Kühl

## Abstract

The internal light field and thus light exposure of the photosymbiotic microalgae (*Symbiodinium* sp.) in corals is strongly modulated by the optical properties of coral tissue and skeleton. While there are numerous studies documenting the light microenvironment in corals, there are only few measurements of the inherent optical properties of corals in the literature, and this has hampered a more quantitative understanding of coral optics. Here we present a study of the optical properties of 26 live coral samples, representative of 11 coral species and spanning a variety of morphotypes. We employed well-established fiber-optic reflectance spectroscopy techniques from biomedical optics using two methods: (1) A source and a detection fiber separated by a variable distance measured the lateral spread of light in corals, dominated by the skeleton; (2) A fiber-optic field radiance probe measured the diffuse reflectance from the coral surface, dominated by the living coral tissue. Analysis based on diffusion theory and Monte Carlo simulation yielded estimates of the bulk scattering and absorption coefficients of the coral tissue and skeleton, in the 750-1030 nm wavelength range. Extrapolating into the spectral region of photosynthetically active radiation (PAR, 400-700 nm) allowed estimation of the optical depth of absorption by the main *Symbiodinium* photopigment chlorophyll a. Coral tissue scattering was on average ~1.9x stronger than the scattering of the skeleton, consistent with the model that corals trap photons by high scattering to enhance absorption by algal pigments, while the lower scattering of the skeleton allows spread of light to otherwise shaded coral tissue areas.

## 1 Introduction

Calcifying, symbiont-bearing corals are the key architects and builders of (sub)tropical coral reefs, one of the most diverse marine habitats on Earth. Coral fitness and thus reef-building is largely dependent on the interactions of the cnidarian animal host and its endosymbiotic microalgae belonging to the dinoflagellate genus *Symbiodinium*. An additional feature of corals is their close interaction with associated microbes that form a diverse microbiome (Bourne et al., 2009). While microbes and nighttime feeding on particles and zooplankton provide essential nutrients and some organic carbon, the coral mainly relies on the supply of labile carbon from *Symbiodinium* photosynthesis (Muscatine et al., 1981). By regulating the nutrient supply to the microalgae and ensuring sufficient light and inorganic carbon supply, the coral host keeps its photosymbionts in a state of unbalanced growth stimulating net O_2_ production and the excretion of photosynthates in the form of simple carbohydrates from *Symbiodinium* to the host (Falkowski et al., 1984). It is estimated that *Symbiodinium* photosynthesis can cover up to >90% of the energy demand of the coral host (Muscatine et al., 1981). Optimization of *Symbiodinium* light exposure for photosynthesis and photoprotection against high UV and solar radiation is thus a key trait in corals, and several strategies for such optimization have been identified in corals (summarised in Wangpraseurt, 2016). Coral optics have also been linked to coral bleaching susceptibility (Enriquez et al., 2005; Rodriguez-Roman et al., 2006; Swain et al., 2016; Wangpraseurt et al., 2017a).

The use of fiber-optic microprobes (Kühl et al., 1995; Wangpraseurt et al., 2012) and various types of reflectance measurements (Enriquez et al., 2005; Marcelino et al., 2013; Salih et al., 2000; Wangpraseurt et al., 2014) have revealed the presence of light gradients and intense scattering of light in both coral tissue and skeleton, which are modulated by coral tissue plasticity and thickness (Wangpraseurt et al., 2014), and the presence and distribution of fluorescent and scattering host pigments in the coral tissue (Lyndby et al., 2016; Salih et al., 2000; Smith et al., 2017). Studies of clean coral skeletons indicate that differences in defined skeleton scattering properties can also play an important role for the light field in corals (Marcelino et al., 2013), which was further supported by Enríquez et al. (2017) studying different museum coral skeleton specimens for relative light enhancement, a less defined measure of scattering.

Modelling and simulation of coral light fields based on estimates of inherent optical parameters remain underexplored with so far only 2 publications in the literature (Teran et al., 2010; Wangpraseurt, 2016). This is mainly reflecting the limited knowledge about the inherent optical properties of corals, i. e., the absorption and scattering coefficients and the angular characteristics of scattering in tissue and skeleton. Recently, we have started to apply experimental methods from biomedical optics such as diffuse reflectance spectroscopy and optical coherence tomography in combination with diffusion theory and Monte Carlo modelling to alleviate this knowledge gap (Wangpraseurt et al. 2016, 2017). In the present study, we present a survey of the absorption and scattering characteristics of 11 different coral species spanning a range of morphotypes. The aim was to obtain robust average values for inherent optical properties of tissue and skeleton of intact corals rather than present detailed high-resolution studies linking such measurements to the fine-structure of individual coral species, which is possible with e.g. optical coherence tomography (Levitz et al., 2004).

## 2 Methods

### Field site and coral sampling

Corals were collected by snorkeling from the reef flat on Heron Island, Great Barrier Reef, Australia (152°06′E, 20°29′S). Corals were fragmented by hammer and chisel and mounted with clay on small tiles for handling and species identification according to Veron (2000). We used a total of 26 fragments representing 11 different species (Supplementary Table 1). Prior to measurements, coral fragments were kept in a shaded outdoor tank at Heron Island Research Station, which was constantly flushed with seawater from the lagoon. At the end of the 5-day measuring period, some fragments were used for determination of *Symbiodinium* cell counts, while the remaining fragments were returned to their original habitat.

### Experimental measurements

Optical fiber-based spectroscopy was used to measure the spreading and reflectance of light from the corals by two different experimental approaches (Figure 1). First, the spreading of light in the skeleton was characterized. Such a point-spread function of light was measured by injecting light into randomly chosen coral tissue areas with one optical fiber collecting light at a distance, r [mm] from the source fiber, which was positioned into the coral tissue (Figure 1A). We note that this volumetric approach mainly probes skeleton scattering. These experiments used 400 μm wide, flat cut optical glass fibers (Ocean Optics, USA), where the source fiber was connected to a white tungsten-halogen light source (HL2000, Ocean Optics, USA), and the collecting fiber was connected to a sensitive fiber-optic spectrometer (QE65000, Ocean Optics, USA). Spectral measurements of light at defined distances from the source fiber, M(r) [counts], were acquired over a wavelength range of 300-1100 nm, where spectral data from 500 to 1030 nm were of sufficient quality for further analysis. Fiber placement was controlled by a manually operated micromanipulator (MM33, Märtzhäuser, Germany), such that the inter-fiber distance, r, was varied in 1 mm steps from 1-10 mm distance from the incident light spot on the coral.

**Figure 1.**
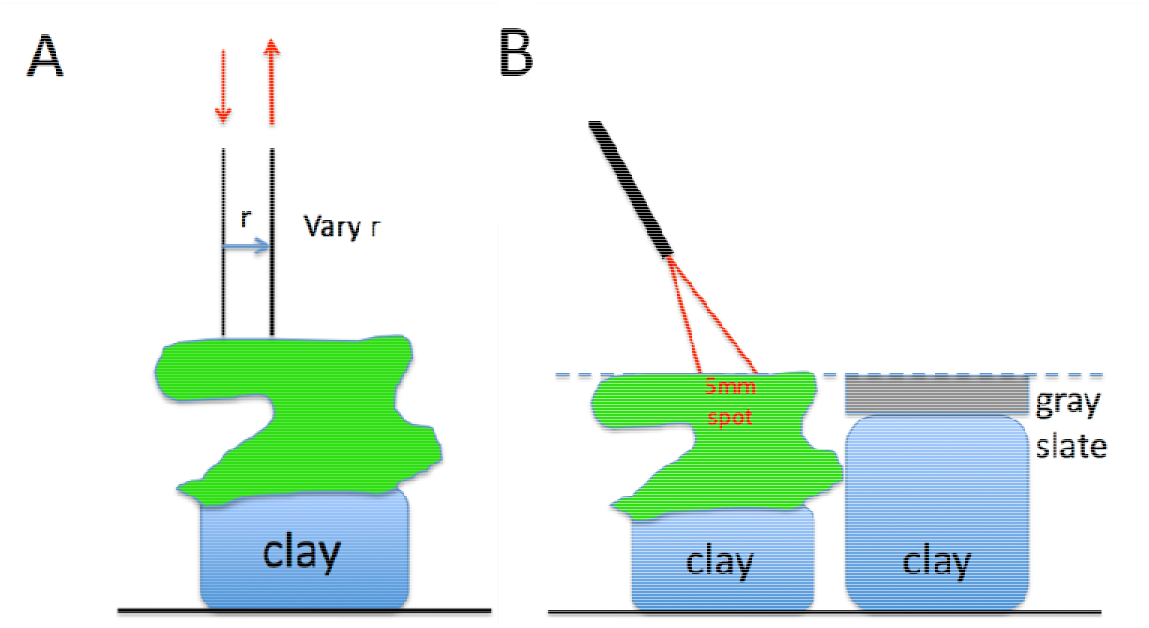
(A) Two optical fibers measure the lateral spread of light within the coral. (B) An optical fiber probe delivers to and collects from a 5-mm-dia. spot on the coral, at a 30° angle off the normal to the coral surface.

Secondly, we measured the diffuse reflectivity of coral tissue, M_d, coral_, relative to the reflectivity of a gray slate reflectance standard, M_d, std_ positioned at identical distance as the coral surface. These measurements were done with a fiber-optic reflection probe (MODEL QR450-7-XSR, Ocean Optics, USA) consisting of 6 illumination fibers surrounding a central collection fiber (all fibers had a diameter of 400 μm). The collection fiber was connected to a fiber-optic spectrometer (QE65000, Ocena Optics), while the illumination fibers were connected to a fiber-optic halogen light source (QE65000, Ocean Optics). The fiber-probe was oriented at an angle of 30° relative to normal with the probe tip positioned at a distance of ~1.5 cm from the coral or reflectance standard surface. This orientation produced a ~5 mm wide spot of illumination on the coral or gray slate standard. The probe collection matched the same 5 mm spot as the light delivery. The 30° illumination mitigated any specular reflectance from the coral or reflectance standard. The coral samples and the reflectance standard were mounted on lumps of clay enabling careful adjustment of their respective heights to ensure identical distance to the fiber probe. After measuring the coral reflectivity, the reflectance standard was shifted into position for measurements allowing correction for any small differences in coral distance from the fiber probe. The reflectance spectrum of a coral was calculated as:

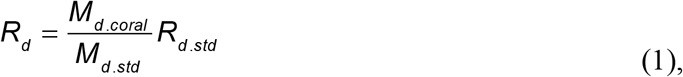

where R_d,std_ is the known reflectance of the reflectance standard (=99%),and M_d,coral_ and M_d,std_ denote the spectral reflectivity of the coral and reflectance standard, respectively.

### Model development

#### Optical properties of skeleton

The measurements of M(r) described the lateral spreading of light within the coral, and hence were dominated by the skeleton optical properties. Such measurements yielded spectra of laterally transported light as a function of radial distance from the source, M(r) [counts], which could be fitted with an exponential function as:

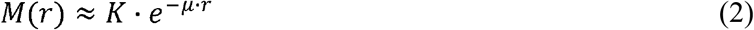

The approximation K·e^−μ·r^, where K is a scaling constant and μ is the attenuation coefficient of light [mm^−1^], ignores the initial 1/r dependence of M(r) at short r, which is expected from diffusion theory (Farrell et al., 1992), since experiments showed that all coral species followed this approximation with no obvious 1/r behavior. Hence, the point-spread function could be well described by the lateral attenuation coefficient μ [mm^−1^], and analysis of M(r) spectra thus yielded values of μ(λ) according to Eq. 2. Figure 2A shows an example of M(r) for 800 nm light, indicating a value of μ = 0.60±0.19 mm^−1^ (n= 26 corals; mean ± SD). Figure 2B shows spectra of μ(λ) for the 26 coral samples, illustrating significant variability among the investigated corals around the overall mean attenuation spectrum (blue line in Fig. 2B).

**Figure 2.**
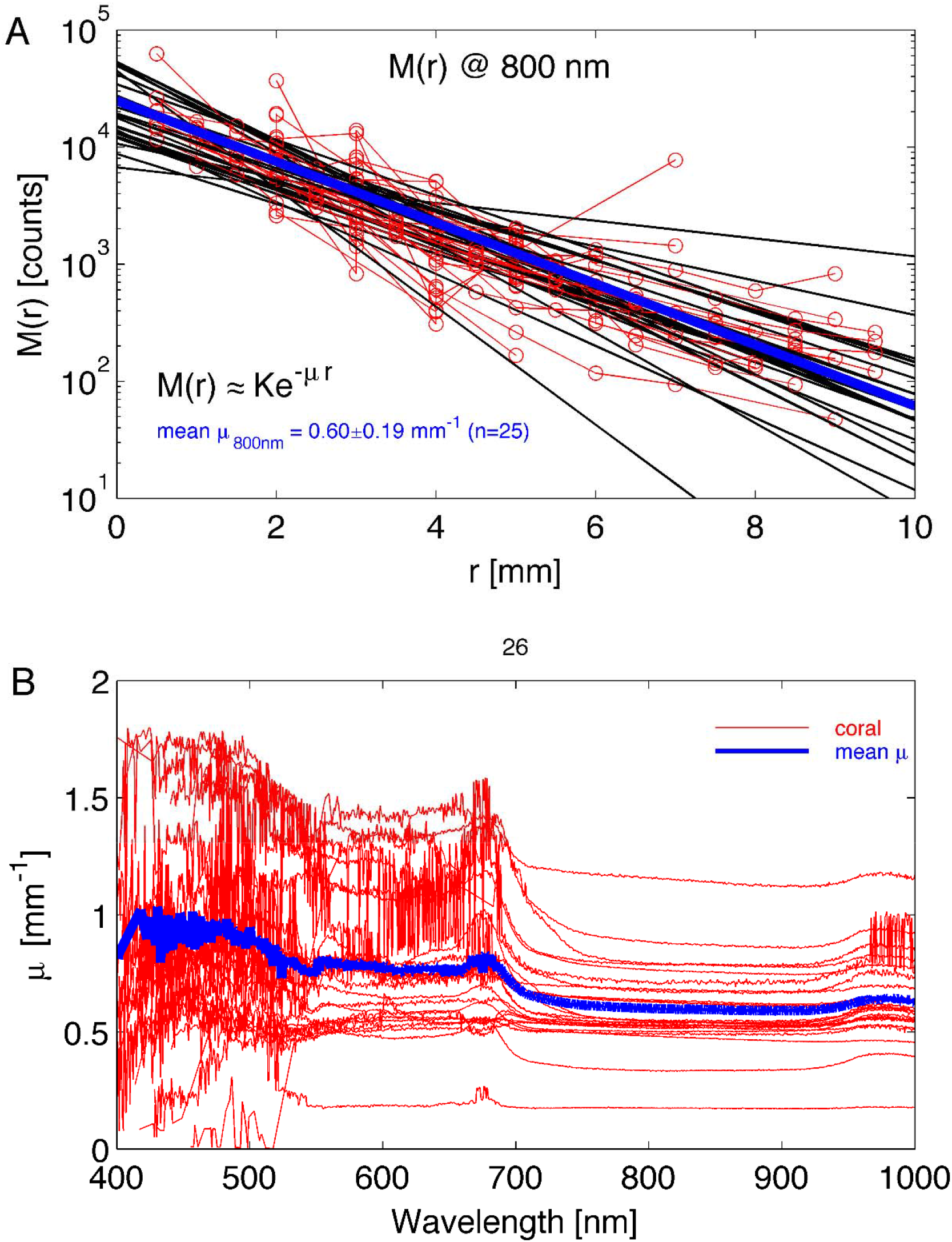
Experimental determination of lateral light attenuation in corals. (A) M(r) data at 800 nm for 26 coral samples fitted with the exponential function M(r) = K·e^−μ·r^ (black lines). Blue line is the average fit. (B) The lateral attenuation spectrum of μ versus wavelength for the 26 coral specimens. Blue line is average spectrum.

Subsequently, μ(λ) was fitted by a least-squares method (multidimensional unconstrained nonlinear minimization, Nelder-Mead, fminsearch.m in MATLAB) to determine the reduced scattering coefficient of the coral skeleton, 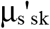 [mm^−1^] and the average skeletal water content, W_s_. The analysis used the 750-1030 nm wavelength range, where algal pigments did not affect the spectrum. The wavelength behavior of 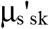 was described by:

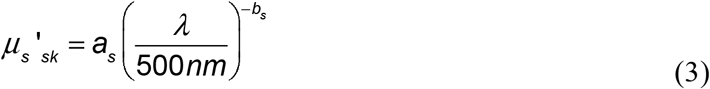

where a_s_ = μ_s_’(500nm), and b_s_ is the skeletal scattering power (Table1).

**Table 1.**
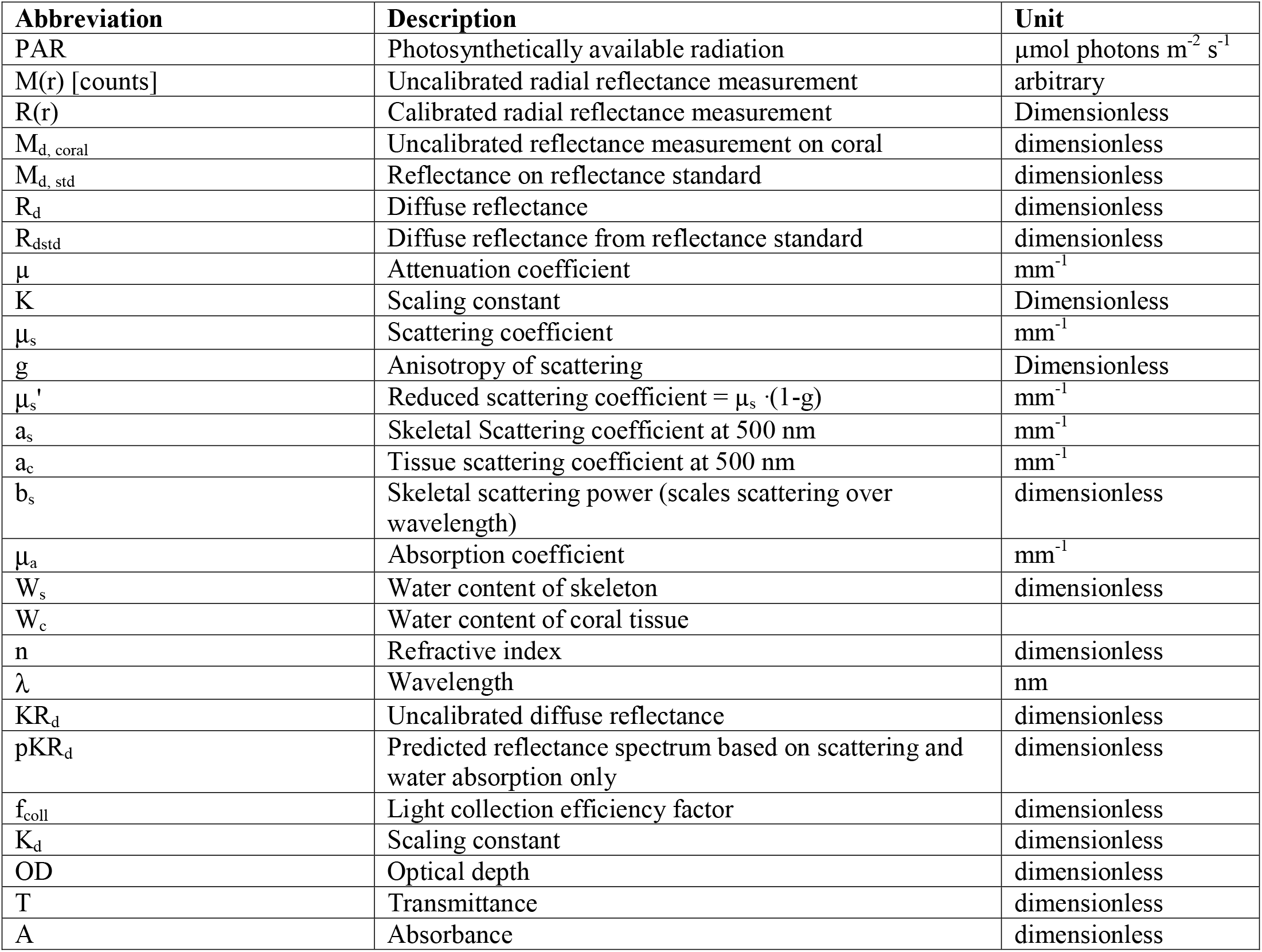
List of abbreviations and their descriptions

Assuming absence of strong spectral signatures by the coral skeleton material (Marcelino et al., 2013), the wavelength dependence of the skeletal absorption coefficient can be approximated by:

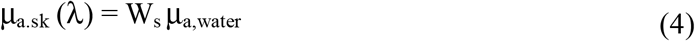

Where W_s_ is the skeletal water content (dimensionless), and μ_a, water_(λ) is the absorption spectrum of water (Hale and Querry, 1973). The analysis used least-squares fitting, where the behavior of the point spread function, R(r),was calculated for each wavelength using a custom written algorithm that calculates the diffuse reflectance based on diffusion theory (Farrell et al. 1992; Tuchin, 2007). This approach has been intensively used in biomedical tissue optics and provides consistent results with other approaches (e.g. Monte Carlo Simulation) when light scattering dominates over light absorption for a given tissue; as a rule of thumb, if μ_s_(1-g)/μ_a_ is >10 (Jacques and Pogue, 2008)). Briefly, the model uses a set of μ_a_ and μ_s_’ values to predict R(r). The model is varied over a range of μ_a_ and μ_s_’ values until a set of values closely match the experimentally measured values. The model assumes a refractive index mismatch between the coral surface and water. We assumed a refractive index of 1.4 for coral tissue and 1.33 for water (Wangpraseurt et al., 2016), thus the refractive index mismatch was calculated as n_r_ = n_coral_/n_water_ = 1.4/1.33.

The R(r) was first interpreted using Eq. 2, R(r) = K·e^−μr^, to yield a simple attenuation coefficient (μ). Iteration then adjusted the values of a_s_, b_s_ and W_s_ until the predicted μ and measured μ agreed. Figure 3 illustrates the ability of least-squares fitting to find the a_s_ and W_s_ of a coral by showing the relative error in a predicted μ versus the experimentally measured μ as:

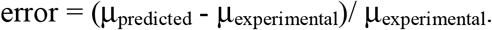

**Figure 3.**
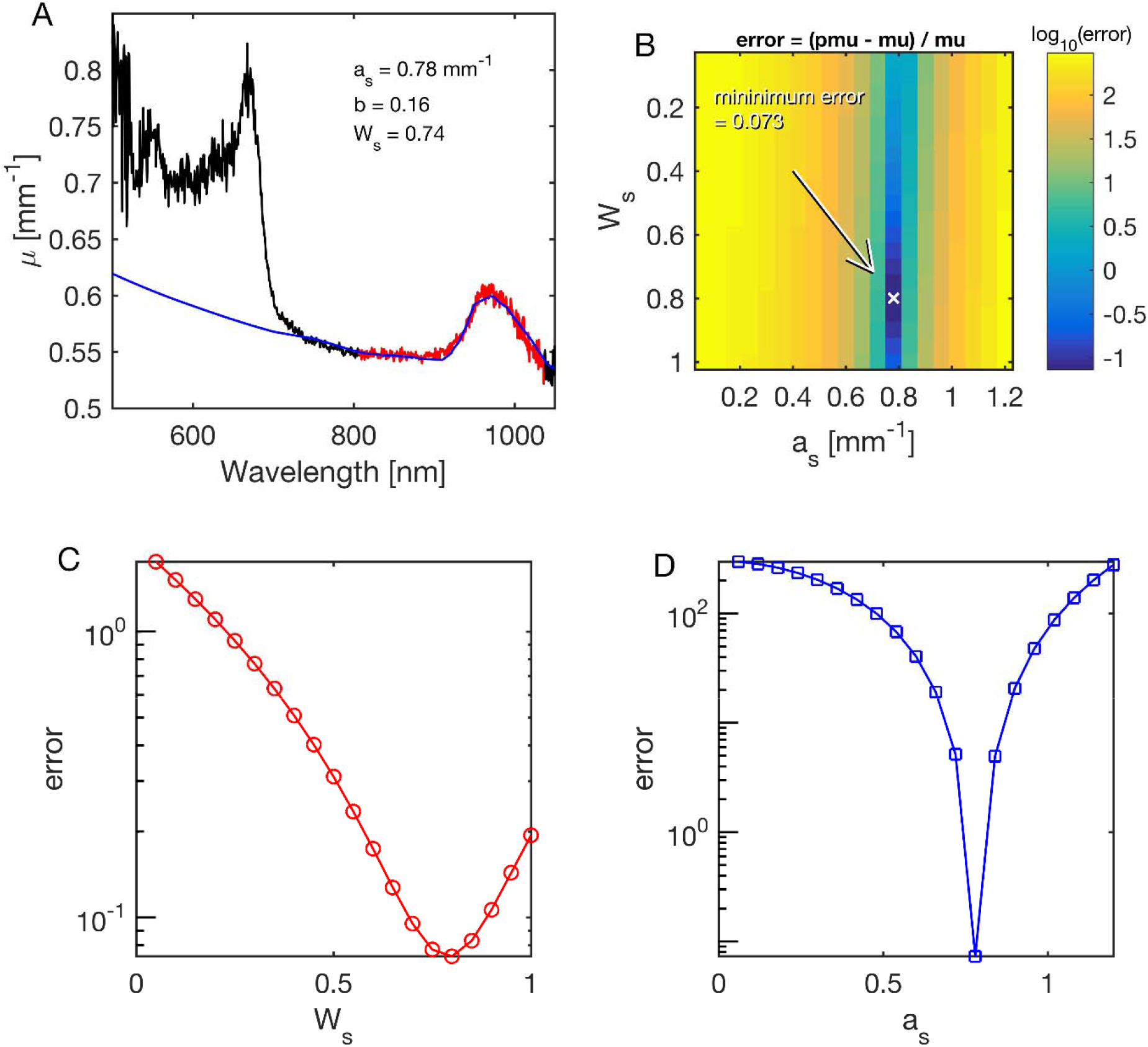
Example of fitting coral skeletal scattering (a_s_) and water content (W_s_) during least-squares fitting of the lateral attenuation coefficient μ(λ) for 750 nm < λ < 1030nm in *Acropora millepora*. The mapping of errors for a_s_ and W_s_, yielded best values of 0.78 mm^−1^ and 0.74 mm^−1^, respectively. (A) The fit (blue line) to the spectral range in red. (B) Map of the error for various choices of a_s_ and W_s_. The parameter b was held at its fitted value of 0.16. Arrow points to minimum error. (C) Error in W_s_, while a_s_ was held at its best value. (D) Error in a_s_, while W_s_ was held at its best value.

#### Optical properties of coral tissue

The diffuse reflectance collected from the 5 mm wide illumination spot was used to estimate the optical properties of the living coral tissue on top of the coral skeleton. The the normalized spectra were divided by the reflectivity of a 99% diffuse reflectance standard (Spectralon) in air, M_d.sp_(λ),to calculate KR_d_ (λ) as:

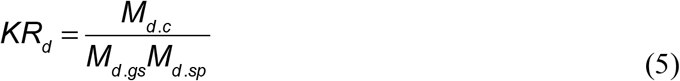

where K is an unknown scaling factor. This procedure canceled the wavelength dependence of the light source and the spectrometer response. Figure 4A shows the raw spectra M_d.c_ (λ), M_d.gs_(λ) and M_d.sp_(λ). Figure 4B shows spectra of KR_d_(λ).

**Figure 4.**
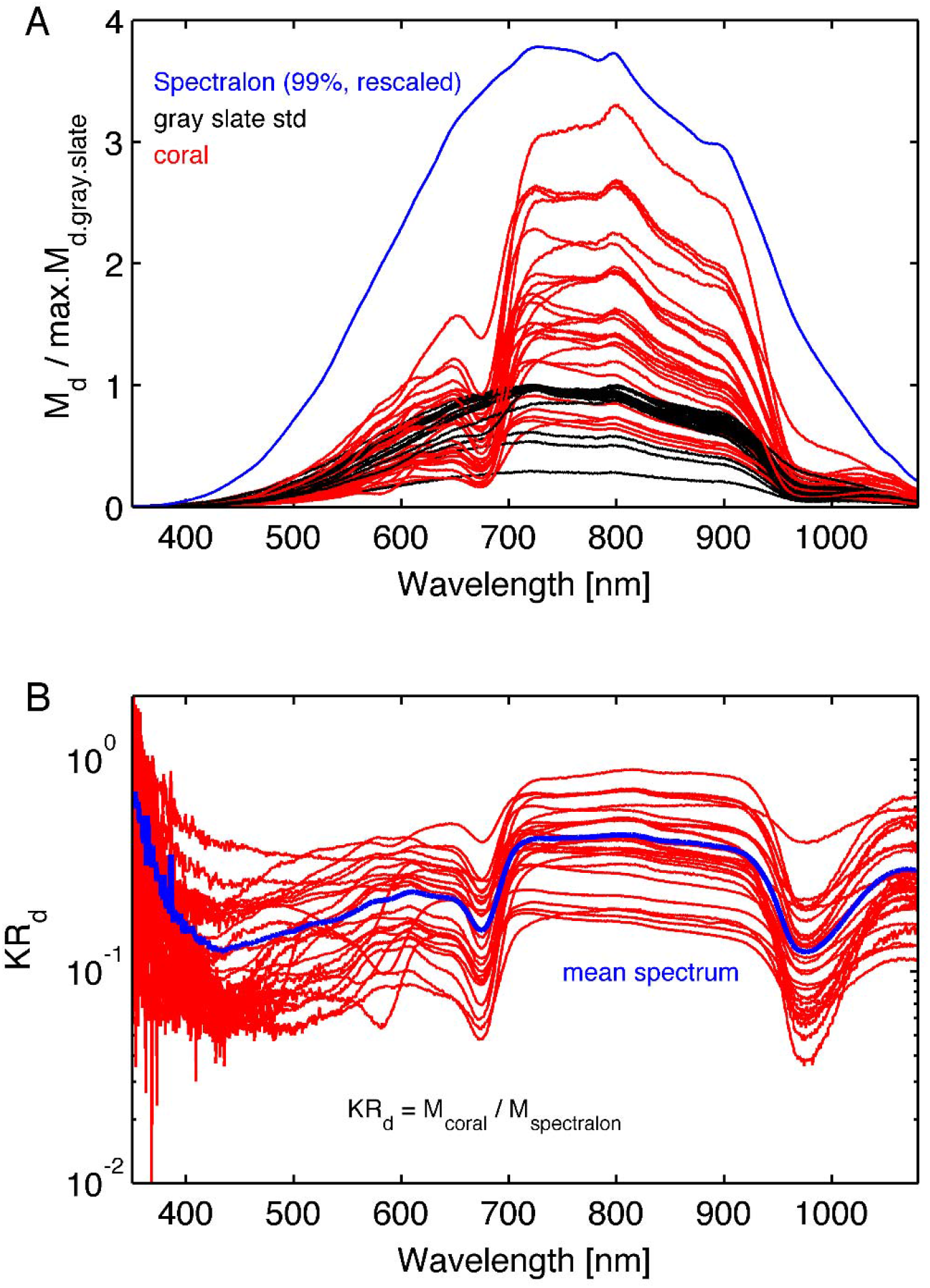
(A) The raw spectra: red lines are the coral spectra M_d.coral_(λ), black lines are reflected light spectra, M_d.gs_(λ), from the gray reflectance standard spectra, and the blue line is reflected light spectrum, M_d.sp_(λ), from the 99% Spectralon reflection standard scaled arbitrarily. The coral and gray reference spectra were normalized by the maximum reflection in the gray reference spectrum for each coral. (B) Normalized reflectance spectra, KR_d_ (λ) (see Eq. 3). The blue line is the mean spectrum of the 26 coral samples.

Each diffuse reflectance spectrum on the coral, KR_d.coral_, was matched by a predicted reflectance spectrum pKR_d_ using least-squares fitting based on the expression:

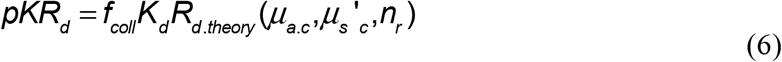

where

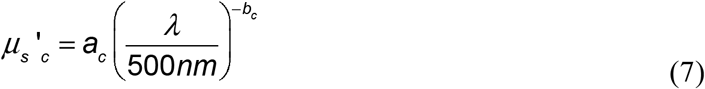

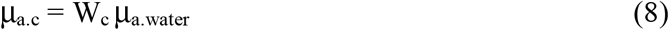

and n_r_ is the refractive index mismatch between coral tissue and water (n_coral_/n_water_ = 1.4/1.33). The fitting parameters were (i) the reduced scattering coefficient at 500 nm, a_c_, and (ii) a scaling constant K_d_.

The coral water content, W_c_, was assumed to be 0.60, which allowed the scattering to be specified so as to match the water absorption peak seen at 960 nm. Such use of water as an internal standard in reflectance spectroscopy depends on the assumed value of the water content, and hence is subject to some, albeit modest, error (Jacques et al. 2010), but the method allows specification of an approximate value for the scattering coefficient. The coral scattering power, b_c_ (Table 1) was assumed to be 0.50, and variation in this assumption had little effect on the fitting.

The factor f_coll_ in Eq. 7 is a light collection efficiency factor specifying the fraction of the total diffuse reflectance collected by the optical fiber probe. The optical fiber probe only collected light from a 5 mm wide spot, and hence failed to collect all the diffuse light escaping the coral. We determined a f_coll_ value of 0.448, by running a two-layer Monte Carlo simulation (Wangpraseurt et al. 2016) that placed a 2-mm-thick living coral layer on top of a semi-infinite skeleton (see more details in the Supplementary Information).

Figure 5 shows an example fit for one of the investigated corals (*Acropora millepora*), where Figure 5A shows the map of errors indicating the locus of minima (white line), while Figure 5B plots the error along this locus, and a blue diamond indicates the minimum along this locus. Figure 5C plots the measured spectrum KR_d_ (red line) and the predicted pKR_d_ (blue line) based on the 750-1030 nm range, i.e., outside the range of coral pigment absorption.

**Figure 5.**
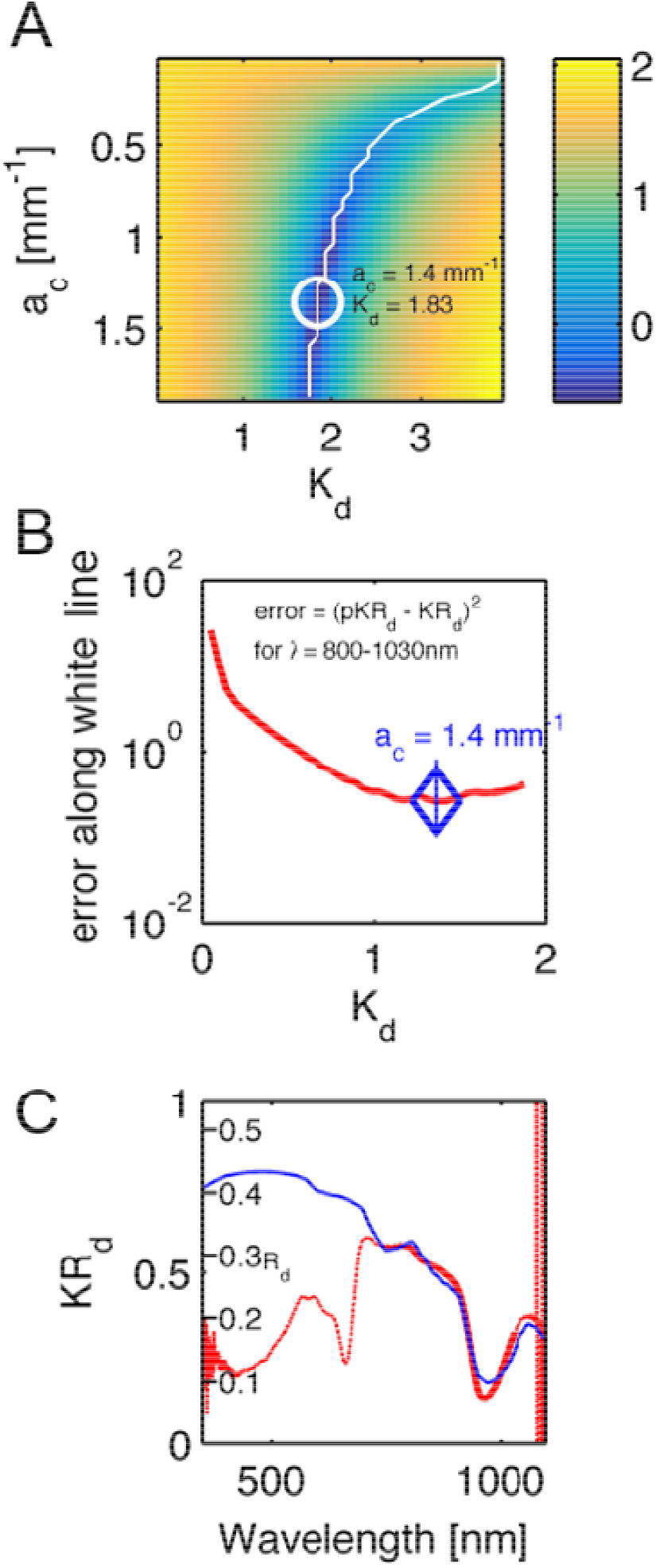
An example fit of the KR_d_ spectrum of a *Acropora millepora*. (A) A map of the errors in measured KR_d_ vs predicted pKR_d_ (Eq. 4), as choices of a_c_ and K_d_ were varied. White line indicates a range of minima. White circle indicates the best minimum using least-squares fitting. Color bar is log_10_ scale. (B) The profile of the error along the white line in Fig. 5A. (C) The experimental spectrum (red line) and the fit to the data from 800-1030 nm. The y-axis is the measured KR_d_. A second set of tic marks labeled Rd indicate the true reflectance collected from the 5-mm-dia. spot by the optical fiber probe, as predicted by R_d_ = pKR/K_d_.

### Algal pigments

The coral spectra shown in Figs. 2 and 4B clearly have absorption by algal pigments at wavelengths <700 nm, including a clear peak absorption in the red part of the spectrum due to Chl *a*, and a range of other pigments at shorter wavelengths. The optical density (OD) is the negative natural logarithm of transmittance (T) and is thus related to absorbance (A) of a material via OD=*A ln10* (Welch and van Gemert, 2011). We can thus calculate OD as a proxy for spectral pigment absorption as:

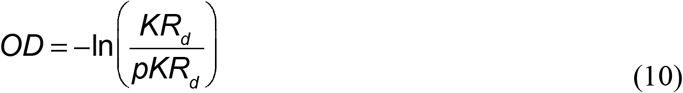

where KR_d_ is the measured reflectance spectrum at the apparent peak of Chl *a* absorption and pKR_d_ is the spectrum devoid of algal pigments as predicted from the scattering plus water absorption (see above). Figure 6 summarizes the values of the OD of Chl *a*, with a mean value of 1.8 ± 0.5 SE.

**Figure 6.**
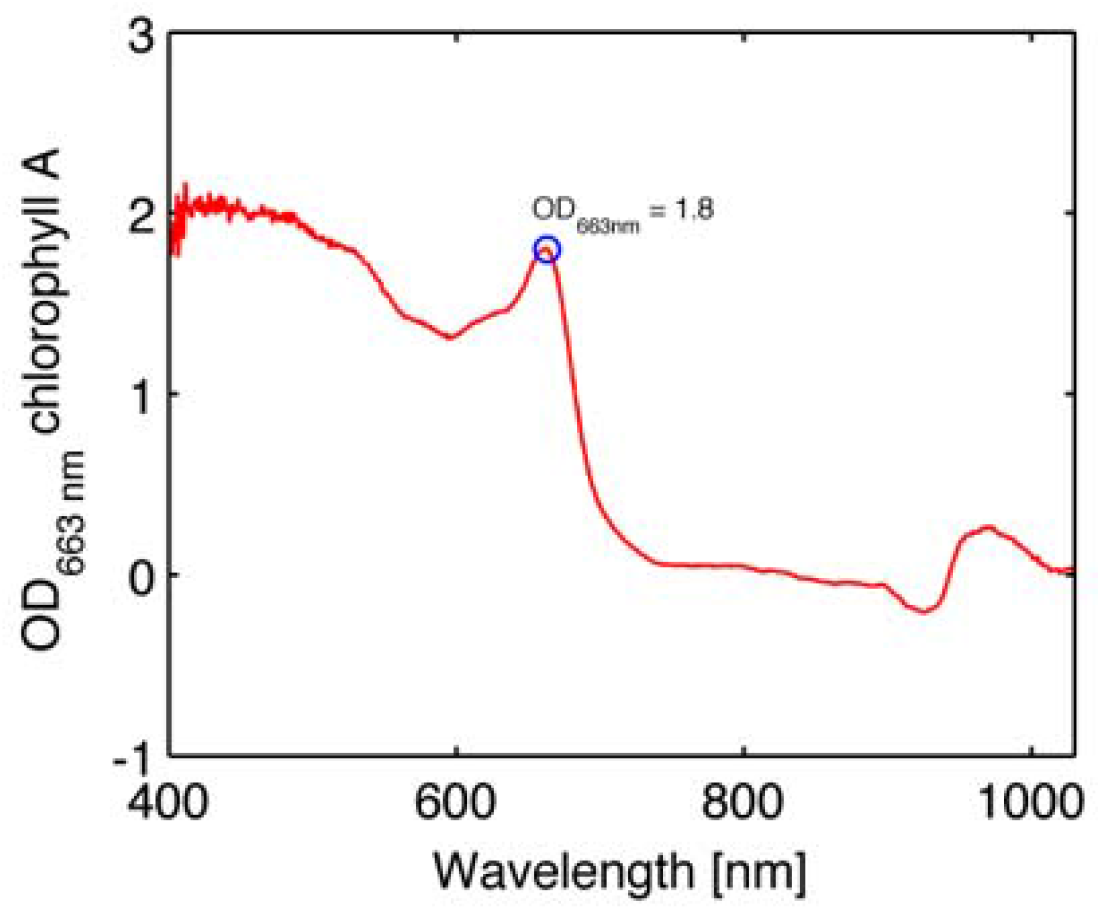
Example of optical density (OD) spectrum of chlorophyll a in the coral *Favites abdita* (Eq. 7).

## 3 Results and Discussion

The present study develops a rapid, non-invasive approach to characterize the optical properties of living intact corals. In contrast to previous studies that focused on analyzing the scattering of dead coral skeletons (Marcelino et al, 2013, Enriquez et al. 2017), we provide a first characterization of optical properties for a range of intact shallow water corals. The main findings of the present study show that coral tissue scattering was ~87% stronger than skeletal scattering for a wide range of investigated corals (Table 2). These results confirm earlier observations of high tissue scattering and low skeletal scattering (Wangpraseurt et al., 2016). It is important to point out what low skeletal scattering implies for coral light transport as different interpretations of the significance of scattering in corals have been promoted (Enríquez et al., 2017; Marcelino et al., 2013; Wangpraseurt et al., 2016).

**Table 2:** Average of coral tissue and skeleton optical parameters

| | $\mu_s'$(500nm) | scatter power | Water content |
| --- | --- | --- | --- |
| | <u>a</u> [$\text{mm}^{-1}$] | <u>b</u> | <u>W</u> |
| Coral skeleton |  |  |  |
| | $0.83 \pm 0.16$ | $0.17 \pm 0.05$ | $0.64 \pm 28$ |
| Coral tissue |  |  |  |
| | $1.55 \pm 1.06$ | (0.5 assumed) | (0.6 assumed) |

Marcelino et al. (2013) characterized the reduced scattering coefficient of coral skeletons and showed that corals with low μ_s_’ are characterized by a higher spread of light within the skeleton. In contrast, Enriquez et al. (2017) visualized the spreading of a laser beam on coral skeletons and argued that high scattering in coral skeletons leads to a high spreading of light. The relationship between light spreading and skeletal scattering is however non-linear and requires detailed quantification of μ_s_’ values. We illustrate this with a simple Monte Carlo model-based simulation of light propagation and fluence rate (= scalar irradiance) in a coral model (see Supplementary information). Assuming a simple light absorbing medium (i.e. μ_s_’=0), an incident collimated light beam will attenuate exponentially according to Lambert Beer’s law. All light will be absorbed along the central line of illumination and no lateral spread of light is visible (Fig. S1a,b). Using a low skeletal scattering value (μ_s_’ = 1 mm^−1^), the fluence rate spreads within the skeleton and can be measured as a high radial reflectance away from the point of illumination (Fig. S1c). In this scenario, (μ_s_’= 1 μ_a_=0.1 mm^−1^) increasing μ_s_’ has increased the spread of light. However, increasing μ_s_’ 10 fold to μ_s_’ = 10 mm^−1^ (e.g. Marcelino et al. 2013), reduces the spread of light (Fig. S1d). Now higher skeletal scattering leads to lower spreading of light, and most of the light is backscattered from a smaller area within the skeleton. Visual observations of intense light spreading of a laser beam from a skeleton do not imply that the scattering coefficient of the skeleton is high (Enriquez et al. 2017).

It has been argued that skeletal light scattering plays a key role in enhancing photosynthetic efficiency of corals (Enriquez et al. 2006), which can be disadvantageous during periods of excess irradiance, where the scattered light causes additional stress leading to photoinhibition and loss of symbionts further increasing light stress in the tissue (Marcelino et al., 2013; Teran et al., 2010; Wangpraseurt et al., 2017a). However, it has also been reported that coral tissue scattering can have a central role in affecting coral light propagation and the contribution of skeletal scattering to coral light absorption (Wangpraseurt et al., 2012; Wangpraseurt, 2016; Wangpraseurt et al., 2014). To illustrate the combined role of tissue and skeletal scattering in coral light transport, we developed a set of Monte Carlo simulations based on the optical properties determined in the present study (see Supplementary information). We first used average optical properties, i.e., high tissue scattering and lower skeletal scattering (Table 2). Secondly, the coral tissue scattering was reduced 10-fold to test whether the coral tissue scattering had an effect on light absorption by Chl *a*. We calculated the intregal of fluence rate in the coral tissue layer, which is proportional to the amount of light absorbed by Chl *a*. The results show that the scattering properties of the living coral tissue clearly affects how light is concentrated in that layer, leading to an approximate 17% increase in light absorption for the high tissue scattering scenario (Fig S2a). With high coral tissue scattering, the photons are trapped in the coral layer and thus enhance the absorption, while low coral tissue scattering leads to photons passing through the coral layer entering the skeleton, where it is backscattered to the coral tissue layer. The oblique angle of backscatter of diffuse light doubles the opportunity of algae to absorb light compared to the nearly collimated incident sunlight that enters the coral (Enriquez et al. 2005; Kühl and Jørgensen 1994).

*Symbiodinium* density controls the absorption of Chl *a*, and *Symbiodnium* cell density is highly variable between corals and over time due to e.g. seasonal fluctuations and/or environmental stress (Hoegh-Guldberg and Jones, 1999). Therefore, we also varied the absorption coefficient over a range of values, (μ_a_= 0.001 to 0.050 mm^−1^) corresponding to a range of realistic Chl *a* concentrations (Teran et al. 2010). Using a high absorption coefficient (μ_a_ =0.050 mm^−1^), our Monte Carlo simulations, show that high coral tissue scattering leads to a 6.8 % decrease in light absorbed by Chl a, relative to the low coral tissue scattering scenario (Fig. S2). Thus, depending on *Symbiodinium* cell density, increasing tissue scattering can increase or decrease light absorption. This is an important finding as it suggests that the role of light scattering in corals has to be evaluated with respect to algal cell density and Chl *a* content (Fig. S2a,b). Additionally, it is important to consider that a realistic prediction of light absorption by *Symbiodinium* cells would also require to take into account the structural complexity of both coral tissue (e.g. tissue thickness, surface structure, *Symbiodinium* distribution, and coral host pigments) (Wangpraseurt et al., 2017b) and skeletons (Enríquez et al., 2017), which is beyond the scope of the present study.

We found no species-specific differences in optical parameters (see Supplementary Table S1, ANOVA p>0.05). Mean tissue scattering [mm^−1^] ranged between 1.0 (Poritidae) - 2.2(Acroporidae) and was not significantly different between the investigated coral families (Figure 7A, ANOVA, p>0.05). Grouping of optical parameters in coral families showed that the average skeletal scattering ranged between 0.7-0.8 [mm^−1^] and was not significantly different between the four coral families (Figure 7A, ANOVA, p>0.05). Our results show that due to the small scattering power of the skeleton (b=0.17), skeletal scattering varies little with wavelength between 400 nm (μ_s_’=0.86) and 800 nm (μ_s_’=0.76). The estimated skeletal scattering is similar to previous estimates from a Faviid coral (about 0.3-0.4 mm^−1^; Wangpraseurt et al. 2016). In contrast, Marcelino et al. (2013) and Swain et al. (2016) have reported species-specific differences in skeletal light scattering and their estimates are about one order of magnitude higher than reported here. However, Marcelino et al. (2013) used low coherence enhanced backscattering spectroscopy to determine the reduced scattering coefficient for a range of coral skeletons for short photon pathlengths (<100 μm). This approach allows for estimating the reduced scattering coefficient in the locally monitored aragonite structure, while the present approach estimates bulk scattering properties of the skeleton, across the coral surface including the structural complexity of the skeleton and its voids (Fig. S3). Thus, the lower values could be explained by the different measuring techniques.

**Figure 7.**
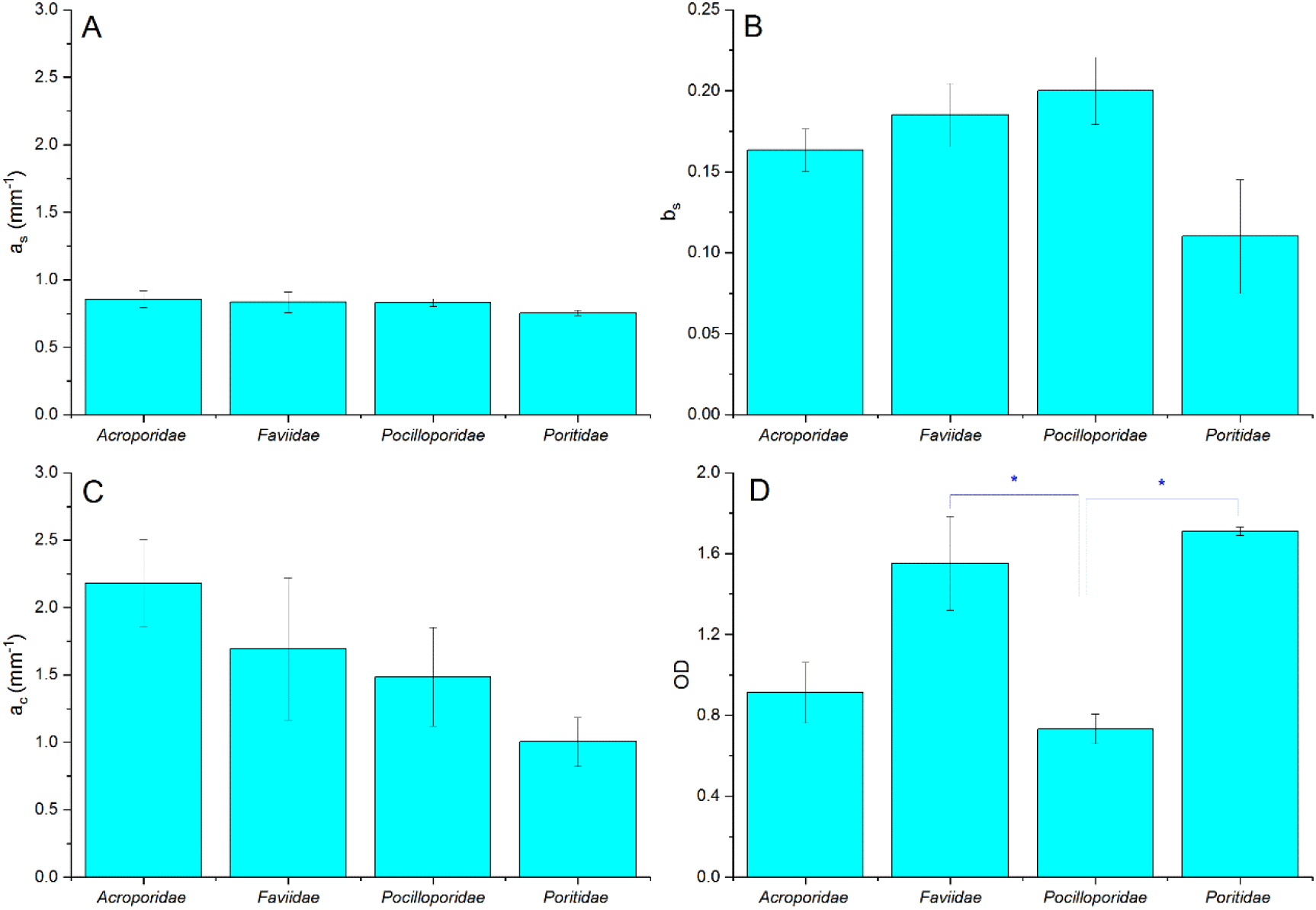
Summary of optical properties of tissue and skeleton for four different coral families. A) skeletal scattering coefficient @ 500nm (a_s_, mm^−1^), B) Skeletal scattering power (b_s_), C) Tissue scattering coefficient @ 500nm (a_c_, mm^−1^), D) Optical depth (OD). Data are means +− SE for *Acroporidae* (n=9), *Faviidae* (n=8), *Pocilloporidae* (n=5) and *Poritidae* (n=3). Signifcant differences (p<0.05) are marked with asterisk.

Although the focus of the present study was on characterizing light scattering, diffuse reflectance spectroscopy also allows for characterizing pigment absorption properties. The optical density (OD) was thus calculated as an indication of pigment absorbance (Figure 7d). We found a mean OD value of 1.8, which characterizes the product of pigment concentration and the average pathlength of photons within the pigmented algal layer. Mean OD differed significantly between coral families, with lowest OD measured in Pocilloporidae (mean =0.73 ±0.07 SE) and highest in Poritidae (mean= 1.71 ± 0.02 SE; ANOVA, p<0.05). In contrast, volumetric cell density estimates suggested highest cell densitites for Pocilloporidae (Supplementary Table 2). The low OD in Pocilloporidae is due to the thin tissue in these corals, as the OD estimates measure OD per projected surface area and thus integrates over the entire tissue volume.

In conclusion, both the coral skeleton and the coral tissue have a central role in modulating light transport and harvesting in corals. The coral skeleton can play a major role in backscattering light toward the overlying living tissue, thereby increasing the harvest of sunlight by Chl a, but the skeleton is translucent and can also play a key role in redistributing light to shaded regions in coral colonies. The efficiency of harvesting sunlight by Chl *a* is affected by high scattering in coral tissue when algal/Chl a levels are low, and diminished when algal/Chl a density is high, which may compensate for fluctuations of low and high algal densities in corals.

## Author contributions

SLJ, DW and MK designed study. SLJ, DW performed experimental measurements. SLJ, DW developed optical simulations and provided analytical tools. SLJ, DW, MK analysed and interpreted data. SLJ, DW, MK wrote the manuscript.

## Acknowledgements

We thank the staff at Heron Island Research Station as well as Sofie Lindegaard Jakobsen, Erik Trampe, Camilla Wentzel and Anthony Larkum for excellent assistance during field work.

## References

Bourne DG, Garren M, Work TM, Rosenberg E, Smith GW, Harvell CD (2009) Microbial disease and the coral holobiont. Trends in microbiology 17: 554–562

Enríquez S, Méndez ER, Hoegh-Guldberg O, Iglesias-Prieto R (2017) Key functional role of the optical properties of coral skeletons in coral ecology and evolution, Proc. R. Soc. B. The Royal Society, p. 20161667

Enriquez S, Mendez ER, Iglesias-Prieto R (2005) Multiple scattering on coral skeletons enhances light absorption by symbiotic algae. Limnology and Oceanography 50: 1025–1032

Falkowski PG, Dubinsky Z, Muscatine L, Porter JW (1984) Light and the bioenergetics of a symbiotic coral. Bioscience 34: 705–709

Farrell TJ, Patterson MS, Wilson B (1992) A diffusion theory model of spatially resolved, steady state diffuse reflectance for the noninvasive determination of tissue optical properties invivo. Medical physics 19: 879–888

Hale GM, Querry MR (1973) Optical constants of water in the 200-nm to 200-μm wavelength region. Applied optics 12: 555–563

Hoegh-Guldberg O, Jones RJ (1999) Photoinhibition and photoprotection in symbiotic dinoflagellates from reef-building corals. Marine Ecology-Progress Series 183: 73–86

Jacques SL, Pogue BW (2008) Tutorial on diffuse light transport. Journal of Biomedical Optics 13: 041302-041302-041319

Kühl M, Cohen Y, Dalsgaard T, Jørgensen BB, Revsbech NP (1995) Microenvironment and photosynthesis of zooxanthellae in scleractinian corals studied with microsensors for O_2_, pH and Light. Marine Ecology-Progress Series 117: 159–172

Levitz D, Thrane L, Frosz M, Andersen P, Andersen C, Andersson-Engels S, Valanciunaite J, Swartling J, Hansen P (2004) Determination of optical scattering properties of highly-scattering media in optical coherence tomography images. Optics express 12: 249–259

Lyndby NH, Kühl M, Wangpraseurt D (2016) Heat generation and light scattering of green fluorescent protein-like pigments in coral tissue. Scientific Reports 6: 26599

Marcelino LA, Westneat MW, Stoyneva V, Henss J, Rogers JD, Radosevich A, Turzhitsky V, Siple M, Fang A, Swain TD (2013) Modulation of Light-Enhancement to Symbiotic Algae by Light-Scattering in Corals and Evolutionary Trends in Bleaching. PLoS ONE 8: e61492

Muscatine L, McCloskey LR, Marian RE (1981) Estimating the daily contribution of carbon from zooxanthellae to coral animal respiration. Limnology and Oceanography 26: 601–611

Rodriguez-Roman A, Hernandez-Pech X, Thome PE, Enriquez S, Iglesias-Prieto R (2006) Photosynthesis and light utilization in the Caribbean coral Montastraea faveolata recovering from a bleaching event. Limnology and Oceanography 51: 2702–2710

Salih A, Larkum A, Cox G, Kühl M, Hoegh-Guldberg O (2000) Fluorescent pigments in corals are photoprotective. Nature 408: 850–853

Smith EG, D’angelo C, Sharon Y, Tchernov D, Wiedenmann J (2017) Acclimatization of symbiotic corals to mesophotic light environments through wavelength transformation by fluorescent protein pigments, Proc. R. Soc. B. The Royal Society, p. 20170320

Swain TD, DuBois E, Gomes A, Stoyneva VP, Radosevich AJ, Henss J, Wagner ME, Derbas J, Grooms HW, Velazquez EM (2016) Skeletal light-scattering accelerates bleaching response in reef-building corals. BMC Ecology 16: 1

Teran E, Mendez ER, Enriquez S, Iglesias-Prieto R (2010) Multiple light scattering and absorption in reef-building corals. Appl Opt 49: 5032–5042

Veron JEN (2000) Corals of the World, vol. 1–3. Australian Institute of Marine Science, Townsville: 295

Wangpraseurt, Larkum AWD, Ralph PJ, Kühl M (2012) Light gradients and optical microniches in coral tissues. Front Microbiol 3

Wangpraseurt D, Holm JB, Larkum AWD, Pernice M, Ralph PJ, Suggett DJ, Kühl M (2017a) In vivo Microscale Measurements of Light and Photosynthesis during Coral Bleaching: Evidence for the Optical Feedback Loop? Frontiers in Microbiology 8

Wangpraseurt D, Jacques S, Petri T, Kuhl M (2016) Monte Carlo modeling of photon propagation reveals highly scattering coral tissue. Frontiers in Plant Science 7:1404

Wangpraseurt D, Jacques SL, Petrie T, Kühl M (2016) Monte Carlo modeling of photon propagation reveals highly scattering coral tissue. Frontiers in plant science 7

Wangpraseurt D, Larkum AWD, Franklin J, Szabo M, Ralph PJ, Kühl M (2014) Lateral light transfer ensures efficient resource distribution in symbiont-bearing corals. Journal of Experimental Biology 217

Wangpraseurt D, Wentzel C, Jacques SL, Wagner M, Kühl M (2017b) In vivo imaging of coral tissue and skeleton with optical coherence tomography. Journal of the Royal Society Interface 14: 20161003

Welch AJ, van Gemert MJ (2011) Optical-thermal response of laser-irradiated tissue. Plenum Press: New York

