## Supplemental file for "Optical properties of living corals determined with diffuse reflectance spectroscopy"

1. **Diffusion theory validation**

For simulating light propagation in corals, a two-layer Monte Carlo method was implemented using the GPU version of MCML (Alerstam et al., 2008; Wang et al., 1995). The optical properties of the skeleton were fixed at the values specified by Eqs. 3 and 4 for light at 800 nm. The optical properties of the 2-mm coral tissue were set to values specified by Eqs. 5 and 6 for 800 nm, after an initial least squares fitting to specify a_c_ and K_d_ while initially assuming f_coll_ = 0.5. The simulation yielded the total diffuse reflectance R_d_ and the point-spread function of escaping photon flux, R (r) [mm^-2^]. This R(r) was interpolated to an R(x,y) spatial distribution, which was convolved against the 5-mm-dia. spot of light delivery to yield the spatial distribution of escaping flux, which was then summed over the area of the 5-mm-dia. collection spot to yield pKR_d_, i.e., the predicted value of KR_d_ based on Monte Carlo simulations. Then, f_coll_ = pKR_d_ /KR_d_. Then the least-squares fitting and Monte Carlo simulation were repeated several times, using the updated f_coll_ each time. The value of f_coll_ stabilized at 0.448. For comparison, the same iterative procedure was applied to a semi-infinite living coral, which yielded f_coll_ equal to 0.450. The factor f_coll_ was not sensitive to whether the living coral tissue was on top of skeleton or semi-infinite living tissue coral. Hence, the use of diffusion theory for analysis was sufficient to characterize the optical properties of the living coral.

2. **Skeletal light spread modeling**

A one-layer Monte Carlo simulation was developed using MCML (Wang et al., 1995). The model was used to exemplify the spread of a collimated light source incident on three planar slabs modeling the coral skeleton with varying scattering strength (Figure S1a-d).

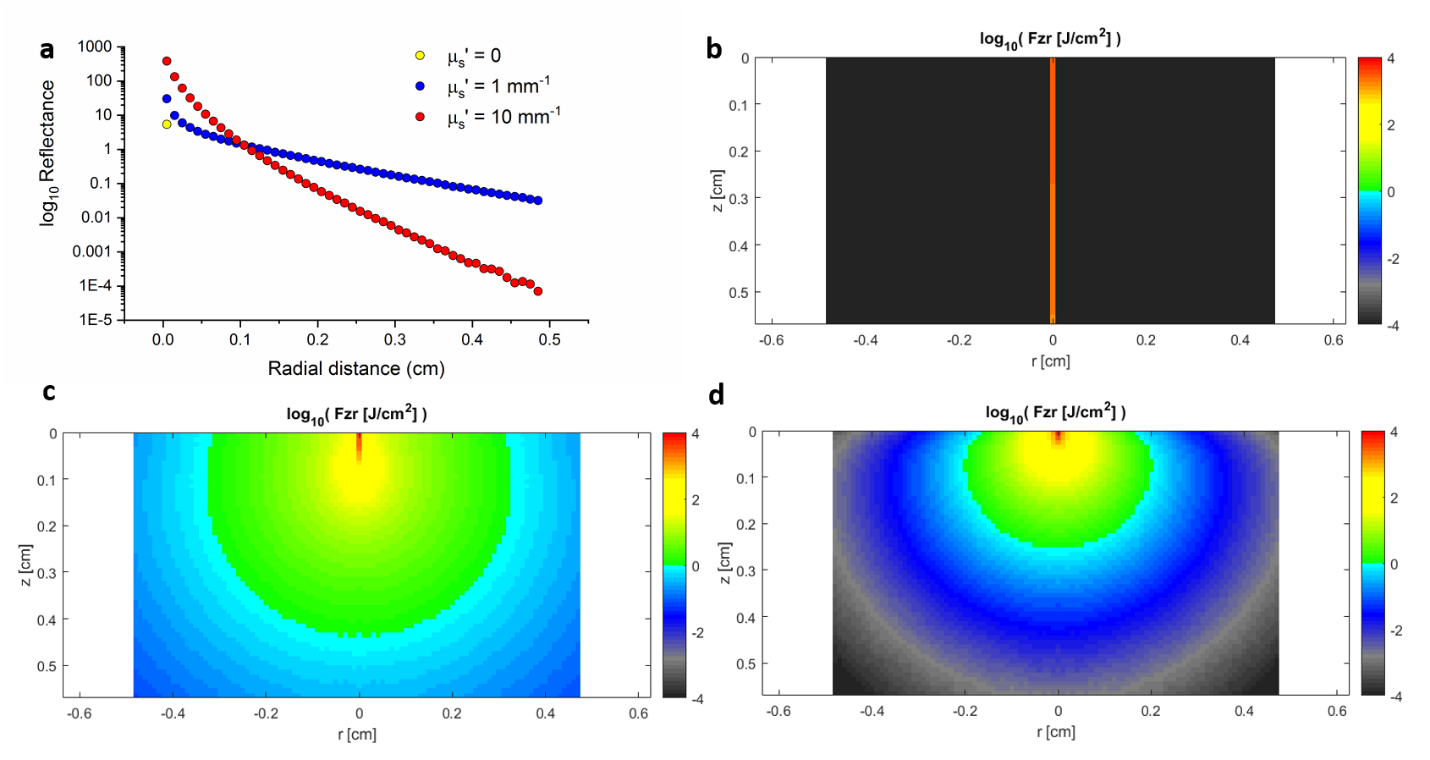

Figure S1 Monte Carlo simulation to illustrate the role of skeletal scattering strength on coral light spreading. The coral skeleton was modeled in planar slab geometry with fixed absorption (µ_a_= 0.1 mm^-1^) and varying scattering strength (µ_s_’=0,1,10 mm^-1^). The lateral escape of light is quantified as radial reflectance (a) and the simulated 2D fluence rate distribution is shown for µ_s_’=0 (b), µ_s_’=1 mm^-1^ (c), µ_s_’=10 mm^-1^ (d).

3. **Example 2-layer model to illustrate light transport in coral tissue and skeleton**

To test the significance of a higher coral tissue scattering versus skeletal scattering, a Monte Carlo simulation modeled light propagation in a 3-mm-thick coral tissue layer over an arbitrarily thick skeleton, where the layers are planar slabs. The 1D axial profiles of fluence rate (=scalar irradiance) (φ, [W cm^2^]) normalized by incident irradiance (E, [W cm^2^]), in other words φ/E [dimensionless], are shown in Fig. S2. The absorption coefficient of the tissue layer was set to a range of values ( µ_a_= 0.001 to 0.050 mm^-1^) corresponding to the absorption at 676 nm wavelength by chlorophyll (Chl) *a* over a range of Chl a concentrations (Teran et al., 2010). The top boundary above the coral was seawater. Two cases were modeled. First, the scattering properties of coral tissue (1.55 mm^-1^) and skeleton (0.83 mm^-1^) at 500 nm wavelength were scaled by µ_s_’ = a·(λ/500)^-1^ for λ = 676 nm to yield values of µ_s_’ = 1.15 mm^-1^ for coral tissue and 0.614 mm^-1^ for skeleton, respectively, at 676 nm. The skeleton absorption was assumed low and identical to the absorption of water, i.e., µ_a_ ~ 4.2x10^-4^ mm^-1^(Jacques, 2013). Secondly, the coral tissue scattering was reduced 10-fold to 0.115 mm^-1^ to test, whether the coral tissue scattering had an effect on absorption of light by coral photosymbionts containing Chl a.

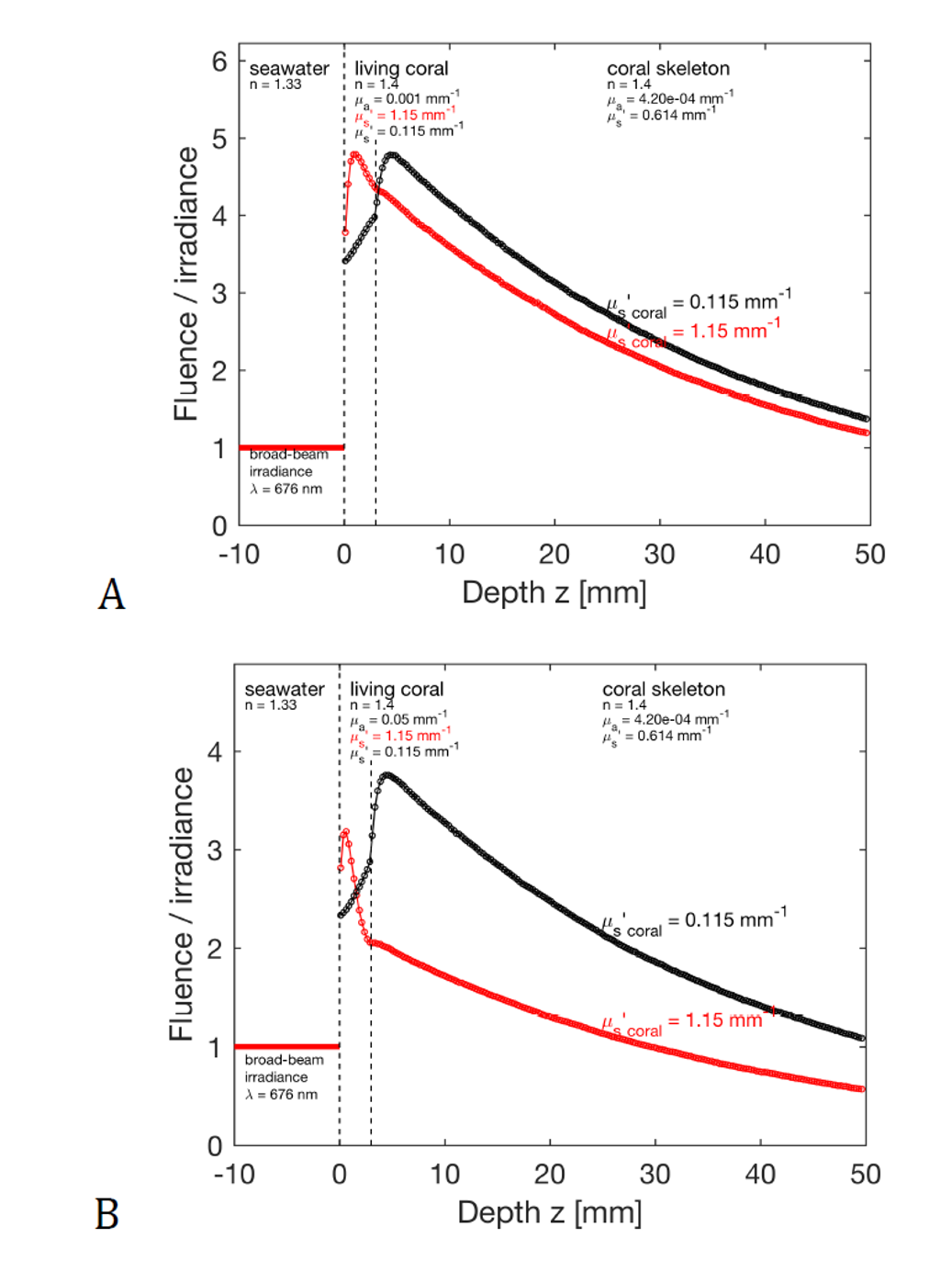

Fig. S2: Simulation of light propagation (676 nm) in corals under uniform illumination for different scattering and absorption scenarios, showing axial profile of fluence rate (= scalar irradiance) normalized to incident irradiance versus depth, for a flat two-layer model of coral (3 mm thick coral tissue layer over a thick skeleton). Simulations show profiles for high (red) and low (black) tissue scattering, respectively. (A) Low Chl *a* concentration scenario, µ_a_ (675 nm) = 0.001 mm^-1^. Integration of normalized fluence rate over the whole coral tissue layer equaled 1.19 and 1.02 under high and low scattering, respectively. Tissue scattering caused a 17% increase in light absorbed by Chl *a* , (B) High Chl *a* concentration scenario, µ_a_ = 0.050 mm^-1^. Integration of normalized fluence rate over the whole coral tissue layer equaled 0.550 and 0.590under low and high scatering, respectively. Scattering lead to a 6.9% decrease in light absorbed by Chl *a*.

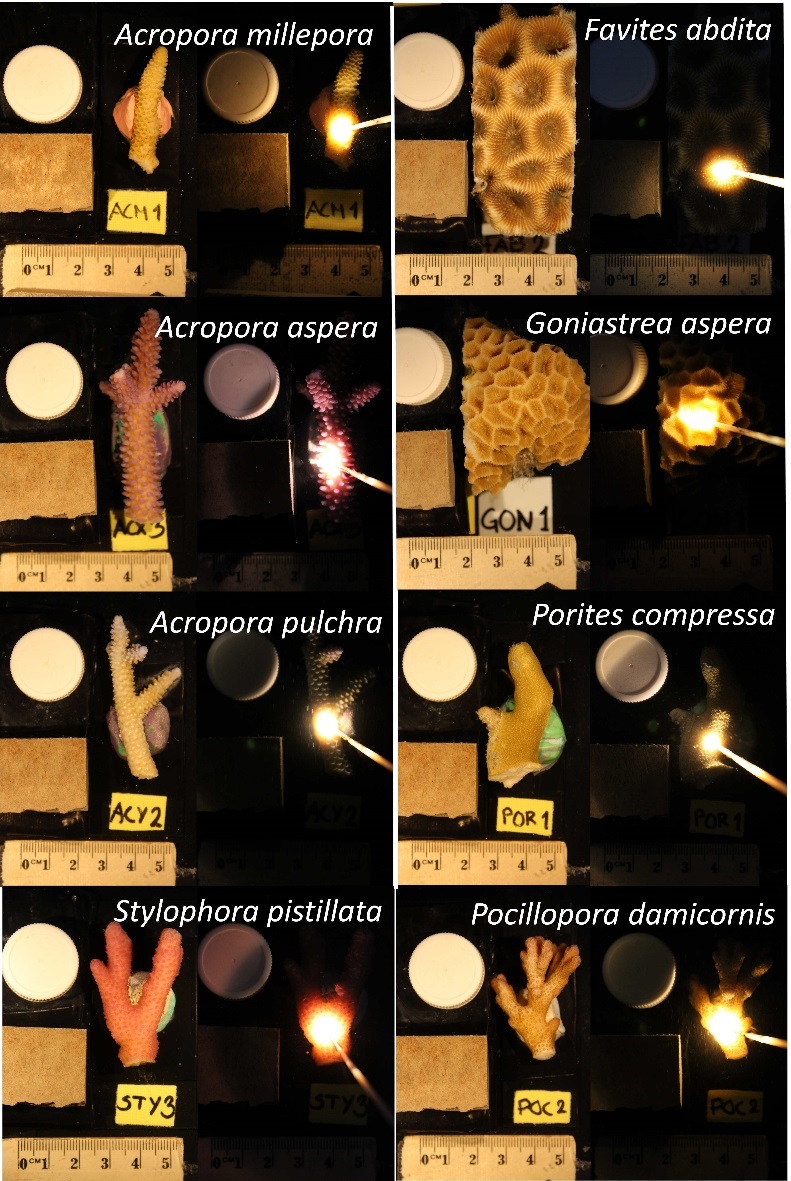

Figure S3. Visualization of light spreading on intact corals. For each species Images were taken on the coral with broadband illumination (left panel) and with an optical fiber (tip diameter= 400 µm) emitting white light at a 45 º angle relative to the coral surface.

Supplementary Table 1: Raw data of extracted optical parameters for all coral specimens.

| **#** | **Species** | **a_s_ mm^-1^** | **W_s_** | **b_s_** | **a_c_**  **mm^-1^** | **K_d_** | **OD** |
| --- | --- | --- | --- | --- | --- | --- | --- |
| 1 | *Acropora millepora* | 0.78 | 0.74 | 0.16 | 1.09 | 1.9 | 1.02 |
| 2 | *Acropora millepora* | 0.86 | 0.49 | 0.18 | 3.47 | 0.5 | 0.50 |
| 3 | *Acropora aspera* | 0.73 | 0.45 | 0.13 | 3.35 | 0.63 | 0.28 |
| 4 | *Acropora aspera* | 0.8 | 0.61 | 0.22 | 1.49 | 1.17 | 0.68 |
| 5 | *Favites abdita* | 0.89 | 1 | 0.18 | 1.78 | 0.6 | 1.41 |
| 6 | *Favites abdita* | 0.98 | 1 | 0.17 | 0.76 | 1.29 | 2.17 |
| 7 | *Favites abdita* | 0.41 | 0.15 | 0.11 | 4.73 | 1.01 | 0.42 |
| 8 | *Favites abdita* | 0.9 | 0.83 | 0.11 | 2.9 | 1.46 | 0.95 |
| 9 | *Goniastrea aspera* | 0.63 | 1 | 0.26 | 0.94 | 1.09 | 1.37 |
| 10 | *Pocillopora damicornis (brown)* | 0.88 | 0.34 | 0.14 | 0.73 | 1.52 | 0.77 |
| 11 | *Pocillopora damicornis (brown)* | 0.77 | 0.33 | 0.19 | 1.29 | 1.12 | 0.90 |
| 12 | *Pocillopora damicornis (pink)* | 0.86 | 0.4 | 0.18 | 2.55 | 0.82 | 0.52 |
| 13 | *Favites abdita* | 1.09 | 0.96 | 0.24 | 1.62 | 1.32 | 1.72 |
| 14 | *Favites abdita* | 0.8 | 0.99 | 0.22 | 0.38 | 2.2 | 2.41 |
| 15 | *Favites abdita* | 0.97 | 0.53 | 0.19 | 0.43 | 2.15 | 1.96 |
| 16 | *Stylophora pistillata* | 0.75 | 0.17 | 0.23 | 2.11 | 0.5 | 0.61 |
| 17 | *Stylophora pistillata* | 0.89 | 0.5 | 0.26 | 0.74 | 1.47 | 0.86 |
| 18 | *Acropora pulchra* | 1.29 | 0.65 | 0.23 | 3.27 | 0.54 | 0.79 |
| 19 | *Acropora pulchra* | 0.73 | 0.9 | 0.12 | 1.05 | 2.38 | 1.10 |
| 20 | *Acropora pulchra* | 0.98 | 0.56 | 0.16 | 1.54 | 2.26 | 0.73 |
| 21 | *Acropora millepora* | 0.77 | 0.84 | 0.13 | 2.08 | 1.56 | 1.40 |
| 22 | *Acropora millepora* | 0.76 | 0.43 | 0.14 | 2.27 | 1.12 | 1.71 |
| 23 | *Porites compressa* | 0.73 | 0.37 | 0.07 | 1.29 | 2.92 | 1.69 |
| 24 | *Porites compressa* | 0.74 | 0.44 | 0.08 | 1.05 | 2.45 | 1.75 |
| 25 | *Porites lobata* | 0.79 | 1 | 0.18 | 0.67 | 1.61 | 1.69 |

Supplementary Table 2. *Symbiodinium* cell density per coral tissue biomass (µl^-1^) for three coral families. *n* are coral colony fragment replicates. For each coral colony, cell counts were measured in triplicate counts in a haemocytometer.

| **Family** | **Mean cells (µl^-1^)** | **SD** | ***n*** |
| --- | --- | --- | --- |
| *Pocilloporidae* | 29.7 | 5.9 | 7 |
| *Acroporidae* | 23.1 | 5.2 | 4 |
| *Faviidae* | 21.3 | 4.0 | 7 |

**References:**

Alerstam E, Svensson T, Andersson-Engels S (2008) Parallel computing with graphics processing units for high-speed Monte Carlo simulation of photon migration. *Journal of Biomedical Optics* 13: 060504-060504-060503

Jacques SL (2013) Optical properties of biological tissues: a review. *Physics in medicine and biology* 58: R37

Teran E, Mendez ER, Enriquez S, Iglesias-Prieto R (2010) Multiple light scattering and absorption in reef-building corals. *Appl Opt* 49: 5032-5042

Wang L, Jacques SL, Zheng L (1995) MCML—Monte Carlo modeling of light transport in multi-layered tissues. *Computer Methods and Programs in Biomedicine* 47: 131-146
